# Nitrogen ionome dynamics on leafy vegetables in tropical climate

**DOI:** 10.1101/2024.10.17.618770

**Authors:** Itamar Shenhar, Aravind Harikumar, Miguel Raul Pebes Trujillo, Zhitong Zhao, Qin Lin, Magdiel Inggrid Setyawati, David Tan, Jie He, Ittai Herrmann, Kee Woei Ng, Matan Gavish, Menachem Moshelion

**Affiliations:** School of Materials Science and Engineering, Nanyang Technological University, Singapore; The Robert H Smith Faculty of Agriculture, Food and Environment, Hebrew University of Jerusalem, Israel; Singapore-HUJ Alliance for Research and Enterprise (SHARE), Singapore; Netatech Engineering Pte Ltd, Singapore; National Institute of Education - Natural Sciences & Science Education, Nanyang Technological University, Singapore; Nanyang Environment and Water Research Institute (NEWRI), Singapore; School of Computer Science and Engineering, Hebrew University of Jerusalem, Jerusalem, Israel

**Keywords:** Nitrogen application, plant ionome, nutrient uptake dynamics, plant physiology, sustainable agriculture

## Abstract

Nitrogen is known to be a critical macro-nutrient influencing plant physiology, growth, and mineral composition. In tropical conditions, which are challenging for leafy vegetable farming, the nitrogen delivery effect is unclear. In this study, we aimed to investigate the effect of nitrogen application on key physiological traits and the mineral composition of the plants, the plant ionome. Experiments were conducted under tropical conditions greenhouse with varying levels of nitrogen supply to examine the effect on plant transpiration, yield, use efficiency of water and nitrogen, and nutrient uptake dynamics followed by cross-correlation analysis, trying to understand the physiological behavior-uptake dynamics relationships. The results demonstrated that transpiration, yield and WUE theoretic optimum curve, which peaking in nitrogen concentration of 120 mg/L for Chinese spinach and 200 mg/L for Chinese broccoli. Conversely, NUE reduce significantly with increasing nitrogen delivery which reflected on antagonistic increase of excess nitrogen.

In terms of mineral composition, nitrogen application resulted in an increase I nitrogen content in the plant leaf tissue, while concentration of certain macronutrients and micronutrients were affected, including potassium, phosphorus, calcium, magnesium, iron, zinc, and molybdenum. Part of the minerals exhibited decreasing pattern due to potential competitive uptake mechanism, iron revealed increasing pattern that correlated with nitrogen delivery, and some minerals correlated with the measured physiological parameters. These results underscore the importance of optimizing nitrogen fertilization to balance plant growth, physiological processes, and plant nutrient homeostasis. The study offers valuable insights for sustainable nitrogen management in agricultural systems aimed at maximizing crop yield while maintaining nutritional quality.

## Introduction

Over the past decade, plant ionome have become a significant research scope in plant science. Plant ionome is determined as the composition of seventeen known essential mineral nutrients and trace elements used for plant growth and development. As nutrient availability is highly dependent on the soil physiochemical properties of the soil (Singh et al., 2022; Hartemink and Barrow, 2023).Plants evolved to maintain mineral homeostasis (Williams and Salt, 2009) and allow physiological resilience under nutrient-changing conditions in order to sustain optimal physiological development under deficiency or excess conditions. Recent studies suggest that the plant ionome has a dynamics role in the crosstalk between different mineral uptake pathways (Fan et al., 2021; Kumar et al., 2021). Ionome crosstalk refers to the change of the mineral composition following a change in mineral supplementation, considering how a change in a mineral concentration can affect and interact with different mineral uptake mechanisms. Most of the studies have been done on a scope of two specific minerals on major crop plants or genetic studies that aim to understand the genetic machinery behind two minerals crosstalk (Lin et al., 2013). For example, a wide mineral dynamics study that On Rapeseed (*Brassica napus* L.), revealed eighteen change situations of minerals uptake following specific mineral deficiency were detected (Maillard et al., 2016) reflecting the biochemical homeostasis dynamics within the plant. Although of those studies, the effect on the crop production, quality, and his connection to the ionome crosstalk is unclear.

In changing environments, mineral uptake is known to limit plant growth (Sinclair, 1992). After carbon (C), nitrogen (N) is the most essential element for plants physiological and morphological development (Hawkesford et al., 2012) as it takes a significant role in the synthesis of proteins, nucleic acids, chlorophyll, cell walls, membranes, and many more primary and secondary metabolites (Marschner, 2011; Mu and Chen, 2021a). Hence, N fertilization can significantly affect the plant physiological properties. Recent studies showed that deficiency and excess in nitrogen delivery reduce plant growth and WUE, which followed by reduced in crop yield (Uhart and Andrade, 1995; Niu et al., 2007; Wang et al., 2011; Mu and Chen, 2021b). Despite the physiological effect, the NUE is increasing in deficiency conditions (Ngosong et al., 2019) and reduce as the concentration increases resulting a negative relationship emphasizing the plant adaptation to the changing conditions.

N availability serves as a fundamental macro element that significantly affect the ionome dynamics in the plant tissues. Previous studies showed that NO_3_^-^ signaling can regulate the Phosphate (P) starvation response in *Arabidopsis thaliana* and demonstrate conserved phenomena response in rice (*Oryza sativa)* and Wheat (*Triticum aestivum*) (Medici et al., 2015; Medici et al., 2019a). Furthermore, on bean (*Phaseolus vulgaris* L.) P starvation led to a decrease in inorganic P concentration in the roots, and NO_3_^-^ simultaneously (Gniazdowska and Rychter, 2000). Ammonium (NH_4_^+^) was also shown to enhance P transport in Corn (*Zea mays* L.) roots (Smith and Jackson, 1987). Additionally, NH_4_^+^ and Potassium (K^+^) were reported to have a competitive relationship in Barley (*Hordeum vulgare* L.) and *Arabidopsis thaliana* roots (ten Hoopen et al., 2010), considered to be due to NH_4_^+^ permeability through voltage-dependent K^+^ channels at the root’s plasma membrane (White, 1996). Zinc (Zn) and nitrogen are suggested to share a unique interaction. Cakmak and Marschner, 1990 suggest that Zn deficiency conditions depressed the net uptake of NO_3_^-^ on Cotton (*Gossypium* L.), Sunflower (*Helianthus annuus)*, and Buckwheat (*Fagopyrum esculentum)*. Furthermore, On Rice *(Oryza sativa)*, Zn and N have positive synergetic effects on the root-to-shoot translocation (Ji et al., 2022). However, a more comprehensive perspective study is needed to clarify the effect of N application on crop yield and ionome so that more species-specific nitrogen application decisions can be made to maximize production and quality.

The impotence of N fertilizer resulted in the overuse of N and a highlighted environmental impact on the soil and the groundwater N contamination (Zhao et al., 2012; Yu et al., 2019). Moreover, manufacturing N fertilizer is a heavy energy consumer that uses fossil fuels and contributes to greenhouse gas emissions (Zhang et al., 2019). Thus, reducing fertilizer use to the actual minimal-optimal level aligns with sustainable agriculture objectives, particularly in regions with a pronounceable environmental footprint from agricultural practice.

In tropical climates, the impact of N supply is unclear. Most of the experiments have been done under chambers with controlled ambient conditions, which can alter the results from common farm conditions and not include a deep analysis of the ionome of the plant, considering mainly the growth and yield, which lack understanding of the crop’s nutritional value (Lee et al., 2024). A study should include physiological yield properties and nutritional outcomes to understand the broad effect of N supplementation on the crop level. Additionally, ionome dynamics can help us understand physiological phenomena that can be difficult to define by standard practice.

Hence, the main objective of this study is to reveal the effect of N supply on the physiological, ionome dynamics, and mineral cross-relationship of leafy vegetables in tropical conditions. To do so, we established an experimental platform that delivers eight fertilizer solution treatments that differ in N concentration and monitor the physiological behavior of the plants as well as their ambient environmental conditions throughout the season. In addition, an ionome analysis was done to understand the crosstalk between N supplementation and ionome dynamics. Our hypotheses were: 1. based on the previous work, is that application of N will have a positive relationship with P and a negative relationship with K; 2. Macro-nutrients will be correlated with the plant’s transpiration and growth traits. 3. An optimum curve will be revealed on the physiological parameters. This study will benefit farmers in tropical conditions by maximizing the yield and quality of production and providing crucial insights into the balanced nutrient management necessary for optimal growth and mineral-dense crops.

## Materials and methods

The study was conducted between September 2023 and November 2023 in a commercial farm greenhouse at Oasis Living Lab, Singapore (1°41’36.0”N, 103°71’95.1”E). We used two commercial leafy vegetable cultivars: *Brassica oleracea* (Alboglabra Group; Chinese broccoli) and *Amaranthus tricolor* L. (Chinese spinach; Supplied by Netatech Ltd., Singapore) in two independent experiments (Fig. 1, A-B). The plants were sewed in seedling trays for 3-4 weeks before being transplanted to 4L pots (“18”, Tefen Ltd, Nahsholim, Israel) filled with coarse sand (“F2”, Rock and Sand industries Ltd, Singapore) for the Chinese spinach and coco-peat (“250”, Riococo Ltd, Sri Lanka) for the Chinese broccoli. The plants grew for 4-5 weeks in a semi-controlled ambient condition greenhouse (Netafim Ltd, Israel), including fans, roof vents, and mesh walls for ventilation.

**Fig. 1.**
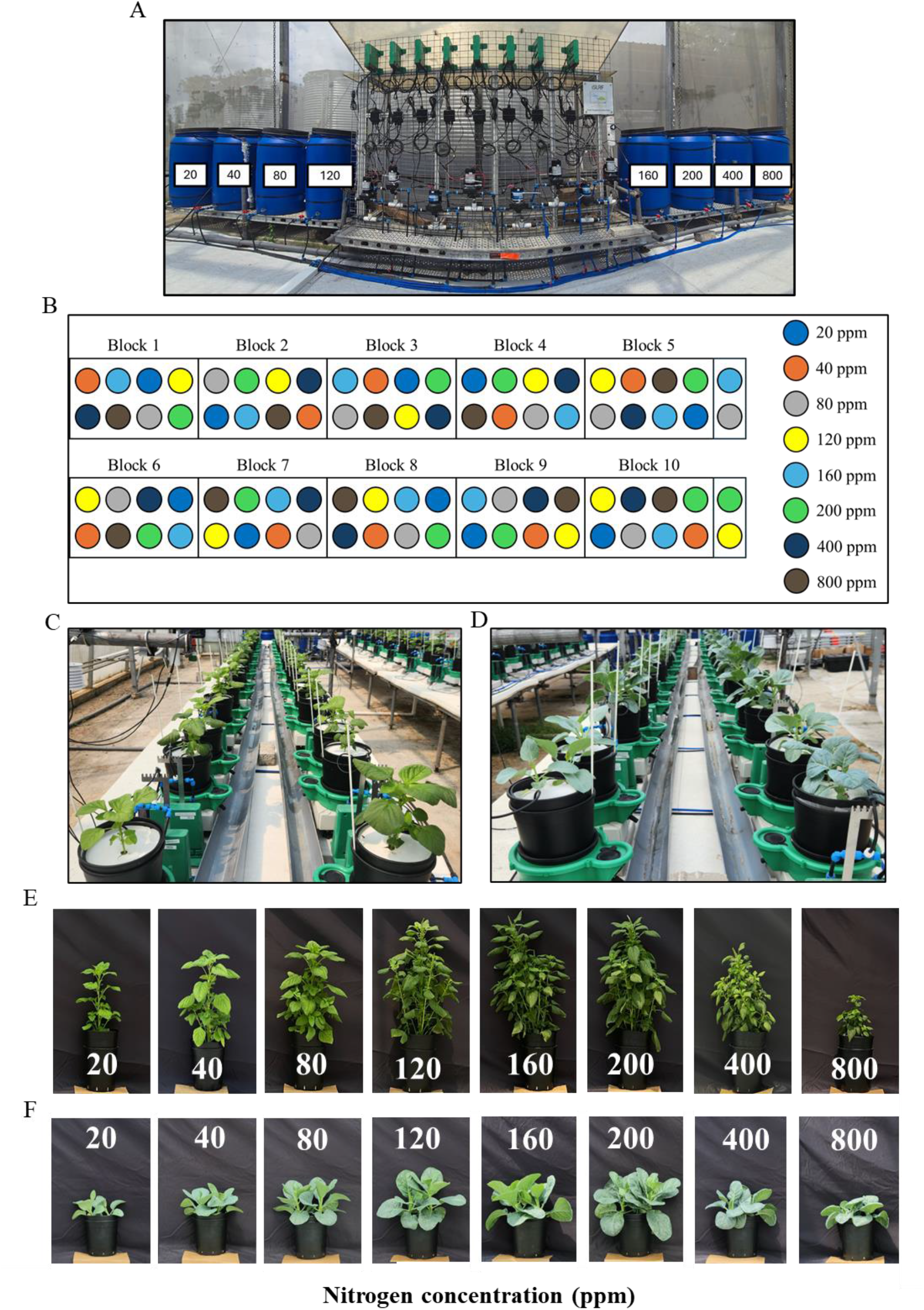
The experimental platform and harvested plants. (A) Panoramic picture of fertilizer solution tanks and automated pump-controller system. (B) Block randomization experiment design of Chinese spinach experiment. Each experimental block includes one repetition from every treatment (C) Picture in the 17 day of the Chinese Spinach experiment. (D) picture on the 27 day of the Chinese Broccoli experiment. Harvest photos of representative plants from the Chinese Spinach (day 45) and Chinese Broccoli (day 30) experiments (E and F respectively).

This study is a spatial and temporal multilevel, multifactorial experiment. Therefore, we used one main meteorological station (“Watchdog 2745”, Spectrum technologies Ltd, Bridgend, Wales) joined with eight smaller meteorological stations (“VP4”, Meter, Washington, USA) to measure the temperature, relative humidity, and vapor pressure deficit (Fig. S1). In addition, we used the main photosynthetic active radiation (PAR) sensor (“#3668”, Spectrum Technologies Ltd, Bridgend, Wales) joined with eight PAR sensors (“SQ-512”, Meter, Washington, United States). Those sensors were connected to the PlantArray system (Plant-Ditech, Yavne, Israel) to maintain a dimensional and continuous measurement. In addition, plants were positioned in a block-randomized design in each experiment to reduce the effect.

Treatments were applied to the plants using pumps (“Shurflo 5050-1311-H011”, Pentair, Minnesota, United States) and controlled by the PlantArray system (Plant-Ditech, Yavne, Israel). Eight different nitrogen concentrations especially suited to the experimental demands were calculated and measured for the study (Table. S1). Continuous EC (“ES-2”, Meter, Washington, United States) and PH (“PHEHT”, Ponsel-Aqualabo, Champigny-sur-Marne, France) were measured during the experiment to monitor the treatment application. The study was conducted on a functional phenotyping platform, PlantArray (Plant-Ditech, Yavne, Israel), to measure continuous physiological parameters. Daily transpiration and cumulative transpiration were measured as described by Halperin et al. 2017 using the experimental protocol described in Dalal et al., 2020. Irrigation was conducted on the plants proportionally according to the measured plant’s transpiration, following four short irrigation events each night to reach soil field capacity and reduce the effect of diverse soil water content on the results.

Plant shoots were harvested and separated manually into leaves and stems. The shoots were dried in a 60°C oven (“DHG-9920A”, Bluepard instrument, Shanghai, China) for one week and measured for dry shoot weight. For mineral analysis, dried leaf tissues of 0.2 g were digested in 4 ml of 70% nitric acid using UltraWAVE single reaction chamber microwave digestion system (Milestone, Sorisole, Italy). The digested solution was diluted with deionized water to a final volume of 25 ml. The Inductively coupled plasma optical emission spectrometer (ICP-OES; “Avio 200”, PerkinElmer Ltd, Massachusetts, United States) and Syngistix software (PerkinElmer Ltd, Massachusetts, United States) were used to measure and calculate the concentrations of minerals (P, K, Ca, Mg, Na, Mn, Cu, Zn, Mo, Fe, B). For the measurement of nitrate concentration, dried leaf samples (0.01 g) were grounded with 10 ml of deionized water and incubated at 37°C for two hours. Sample turbidity was then removed by vacuum filtering the mixture through a 0.45 µm-pore-diameter membrane. The final volume was made to 50 ml. The nitrate concentration of the leaf tissues was determined using the Flow Injection Analyser (Model Quikchem 8000, Lachat Instruments Inc., Milwaukee, USA). Total reduced nitrogen (TRN) content was determined by Kjeldahl digestion of 0.05 g of dried leaf tissues and a Kjeldahl tablet in 5 ml of concentrated sulphuric acid for 60 min at 350°C. After digestion, TRN concentration was quantified by a Kjeltec 8400 analyzer (Foss Tecator AB, Höganäs, Sweden) through titration.

Agronomic transpiration use efficiency (water user efficiency; Agronomic WUE), the ratio between crop yield and transpiration was calculated according to:

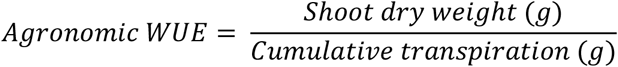

Nitrogen efficiency and balance parameters were calculated according to Congreves et al., 2021:

Nitrogen use efficiency (NUE_crop_) is the ratio of the dry biomass to the supplied N throughout the experiment. It is calculated by:

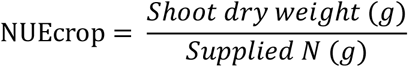

Partial nitrogen balance (PNB) is the ratio of the calculated shoot N to the supplied N throughout the experiment. It is calculated by:

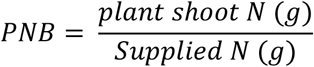

Excess N is the amount of N supplied to the plant and not detected in the plant shoot. It was calculated by:

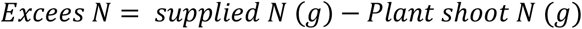

Plant shoot nitrogen was calculated by multiplying the shoot dry weight with the nitrogen concentration.

Data analysis was performed using Microsoft Office 365 Excel (Microsoft Ltd, Washington, United States) and OriginPro 2023 (OriginLab Ltd, Massachusetts, United States) for graph plotting and JMP 17 (SAS Institute Inc., North Carolina, United States) for the statistical analysis. In all multiple comparison tests, data was checked for normal distribution using Shapiro-Wilk’s test and homogeneity of variance using Levene’s test. A Tukey honest significant difference ANOVA test was applied if both tests were satisfied. If the normality or homogeneity of variance criteria were violated, Wilcoxon/Kruskal-Wallis’s nonparametric ANOVA test was used. All tests were done at a p-value<0.05 significance. Data mean values for all the results presented with ± standard error (SE). The squared value of the Pearson correlation coefficient was calculated in a regression fit analysis, and p-values were calculated according to the multivariate coefficient probability test.

## Results

### N alters the physiological behavior of *Amaranthus tricolor L*. and *Brassica oleracea*

Experiments were designed to examine the physiological traits and ionome dynamics of *Amaranthus tricolor L*. (Chinese Spinach) and *Brassica oleracea* (Chinese Broccoli) on different concertation of N. We recognized a significant appearance difference in the other treatments on Chinese Spinach (Fig. 1, C) and Chinese Broccoli (Fig. 1, D). Correspondingly, plants’ daily transpiration throughout the experiments were affected by the N treatments. Treatments 120, 160, and 200ppm showed a significantly higher daily transpiration already 7 days after the treatment start (Fig. 2 A) and maintained this pattern throughout the experiment as revealed in the end of the experiment (Fig. 2B) and the cumulative transpiration (Fig. 2C). Shoot dry weight of these higher transpiring treatments was also significantly higher compared to 20, 40, 80, 400, and 800 treatments (Fig. 2D). An optimum curve was revealed for both cultivars whereas on Chinese spinach 120, 160, and 200 transpired significantly more than other treatments on both cultivars. The plant transpiration-use-efficiency (agronomic WUE, see materials and methods) of treatment range 120-400 exhibited significant higher WUE throughout the experiment duration (Fig. 2E). We further investigate the nitrogen use efficiency (NUE_crop_) and discovered a significant exponential decline pattern of NUE_crop_, where the lower N concentration treatments reveled the highest efficiency (Fig. 2 F). The partial nitrogen balance (PNB) and the excess N revealed NUE_crop_ antagonistic pattern from the PNB (Fig.2, G). Similar results repeated in the Chinese broccoli with the differences on the optimum treatment concentrations of 200ppm exhibited the highest cumulative transpiration and shoot dry weight (Fig. 3, C-D).

**Fig. 2.**
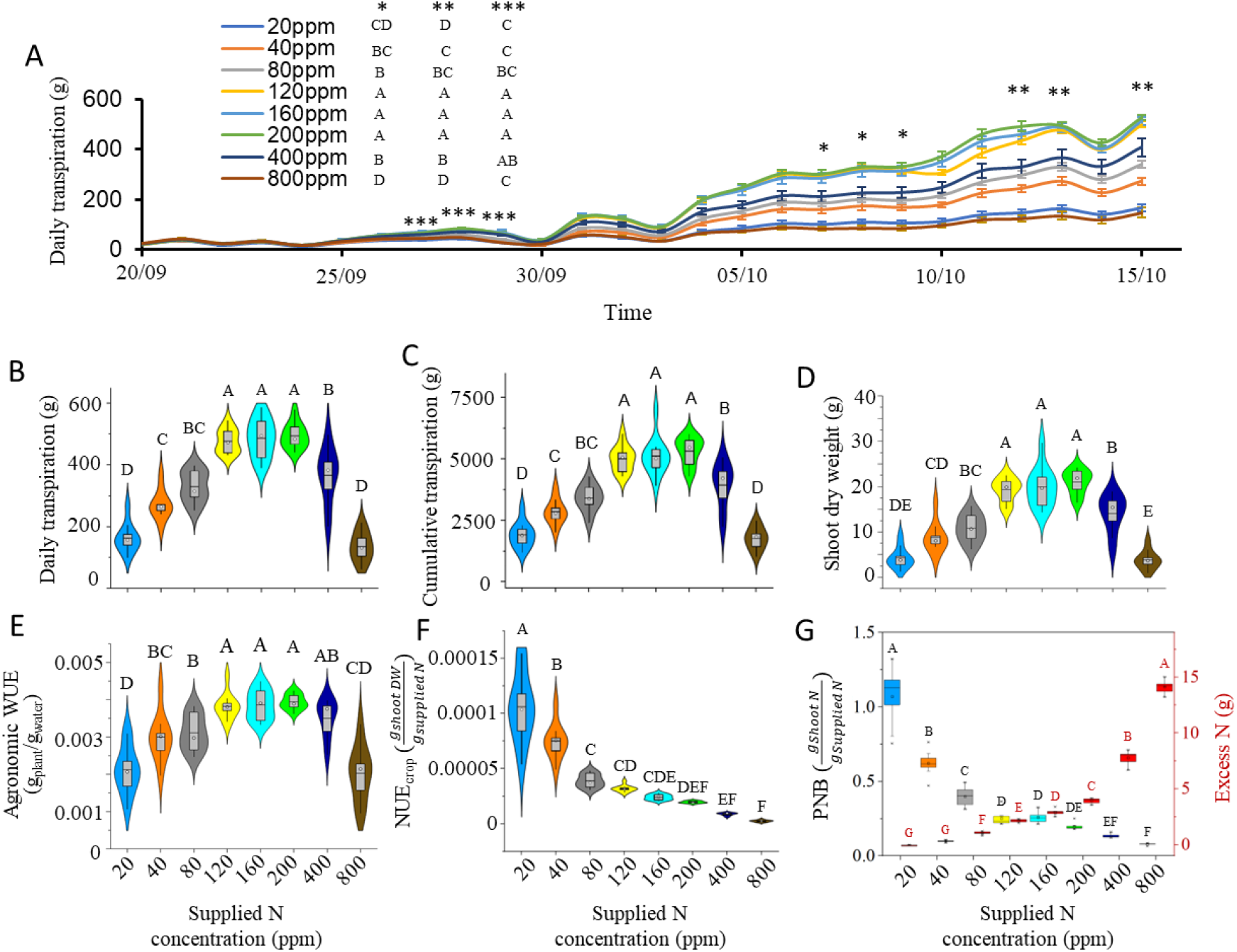
Physiological response of Chinese Spinach to eight nitrogen treatment (A) Daily transpiration 1hmughout the experiment, (B) Daily transpiration on the 13.10.24, (C) Cumulative transpiration all over the experiment, (D) Shoot dry weight, (E) Agronomic WUE, (F) Nitrogen use efficiency (NUE_crop_), and (G) partial nitrogen balance (PNB) and Excess nitrogen during the experiment Different letters represents significant differences using Tukey honest significant difference test (P-value<0.05,9≤N≤11).

**Fig. 3.**
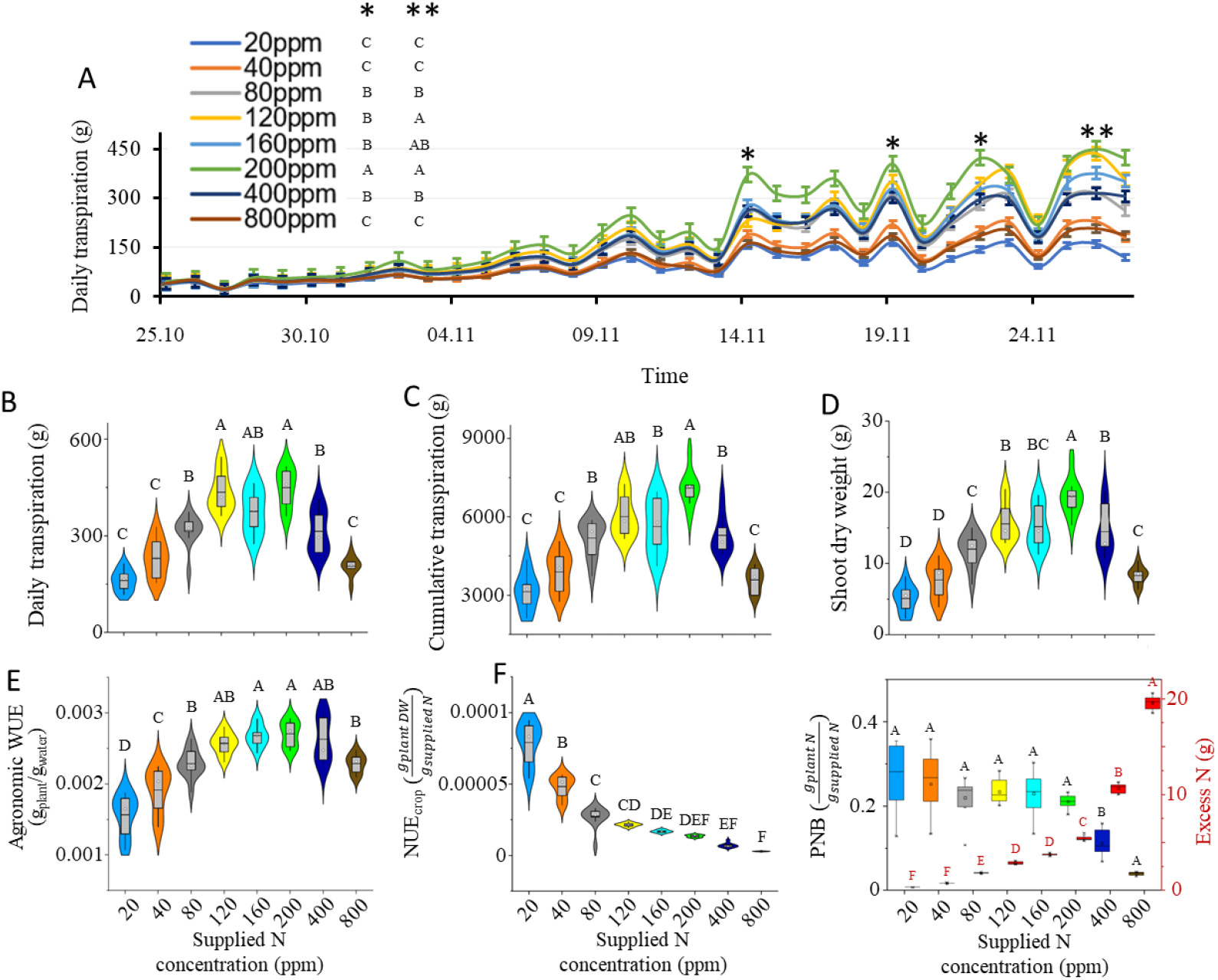
Physiological response of Chinese Broccoli to eight nitrogen treatment. (A) Daily transpiration throughout the experiment. (B) Daily transpiration on the 27-11.24, (C) Cumulative transpiration all over the experiment. (D) Shoot dry weight, (E) Agronomic WUE, (F) Nitrogen use efficiency (NUE_crop_), and (G) Partial nitrogen balance {PNB) and Excess nitrogen during the experiment. Different letters represents significant differences using Tukey honest significant difference test (P-value<0.05, 9≤N≤11)

### N-Ionome dynamics of *Amaranthus tricolor L*. and *Brassica oleracea*

Mineral analysis was preformed to the harvested plant leaf tissue to clarifying the effect of the N treatment on the plant ionome. The NO_3_ and TRN concentrations in the plant tissue exhibited direct correlation with the supplied treatment (Fig. 4, A-B) on both cultivars. This relationship is expected due to the supplied N gradient. Nevertheless, the relationship between the supplied N treatments and other minerals, revealed several different relationship patterns. K showed a negative correlation to N, in particular at the physiological zone (80 – 200 ppm) (Fig. 4 C and D) which reflected on the relationship with leaf N concentration on both cultivars (R^2^=0.787, P-value<0.0001 and R^2^=0.8138, P-value=0.0022; Fig. 6). P concentration revealed non-significant pattern under physiological concentrations of N (80-200ppm; Fig. 4, E and F). Both Ca and Mg displayed similar concentration patterns across N treatments in Chinese spinach and Chinese broccoli. Lower concentrations of both were noted at N concentrations of 20, 40, and 80, with increases observed from 120 to 200. The highest N treatments (400 and 800) significantly reduced concentrations, suggesting a potential toxicity effect. (Fig. 4, G-J). The resembling results between the Ca and the Mg revealed high positive correlation between them on both cultivars (Fig. 6).

**Fig. 4.**
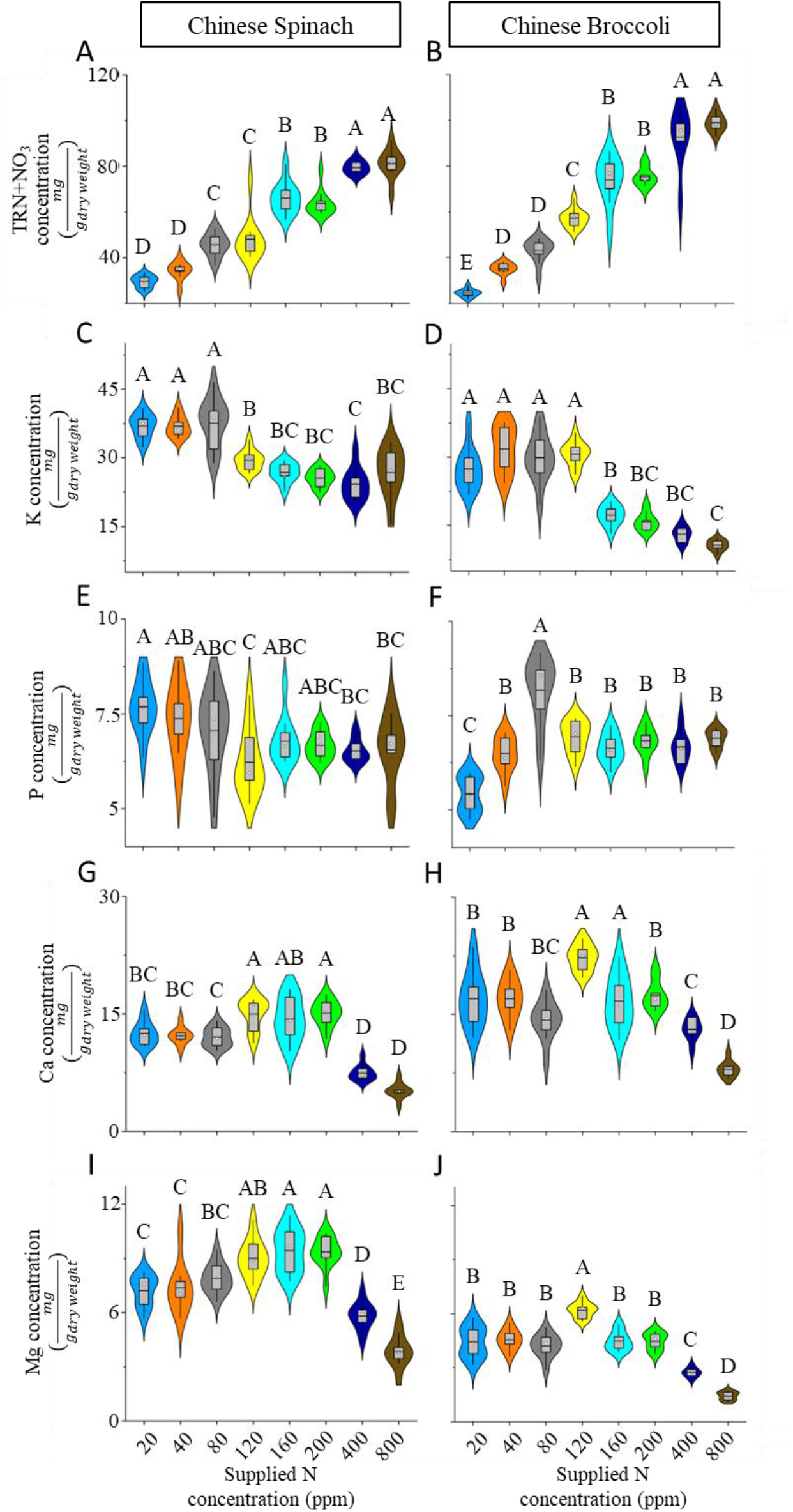
Macro-nutrient effect of the nitrogen treatments of leaf tissue. The macro-nutrients concentrations: (A and B) Nitrogen, (C and D) Potassium, (E and F) Phosphorus, (G and H) Calcium, and (I and J) Magnesium on harvested leaves of Chinese Spinach and Chinese Broccoli respectively. Different letters represents significant differences using Tukey honest significant difference test (P-value<0.05,9≤N≤11).

Fe concentrations in both cultivars increased with N (Fig. 5 A and B), demonstrating strong positive correlations with N (R^2^=0.943 and R^2^=0.84; Fig. 6). The other microelements, Zn, Mn, Cu, and B exhibited non-significant patterns with N at physiological concentrations of 80-200 in both Chinese spinach and Chinese broccoli (Fig. 5, C-J).

**Fig. 5.**
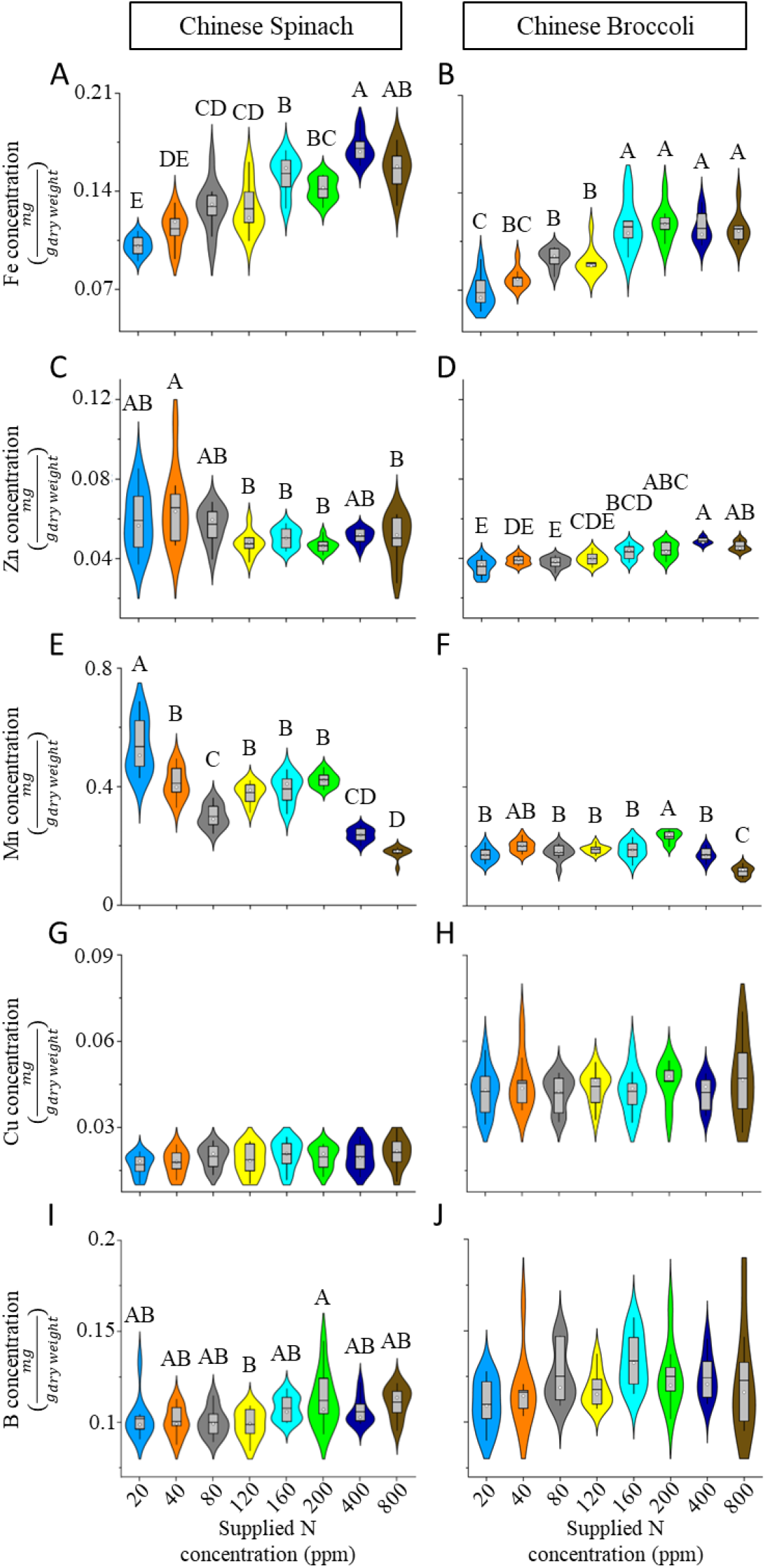
Micro-nutrient effect of the nitrogen treatments of leaf tissue. The macro-nutrients concentrations: (A and B) Iron, (C and D) Zinc, (E and F) Manganese, (G and H) Copper, and (I and J) Boron on harvested leaves of Chinese Spinach and Chinese Broccoli respectively. Different letters represents significant differences using Tukey honest significant difference test (P-value<0.05, 9≤N≤11).

**Fig. 6.**
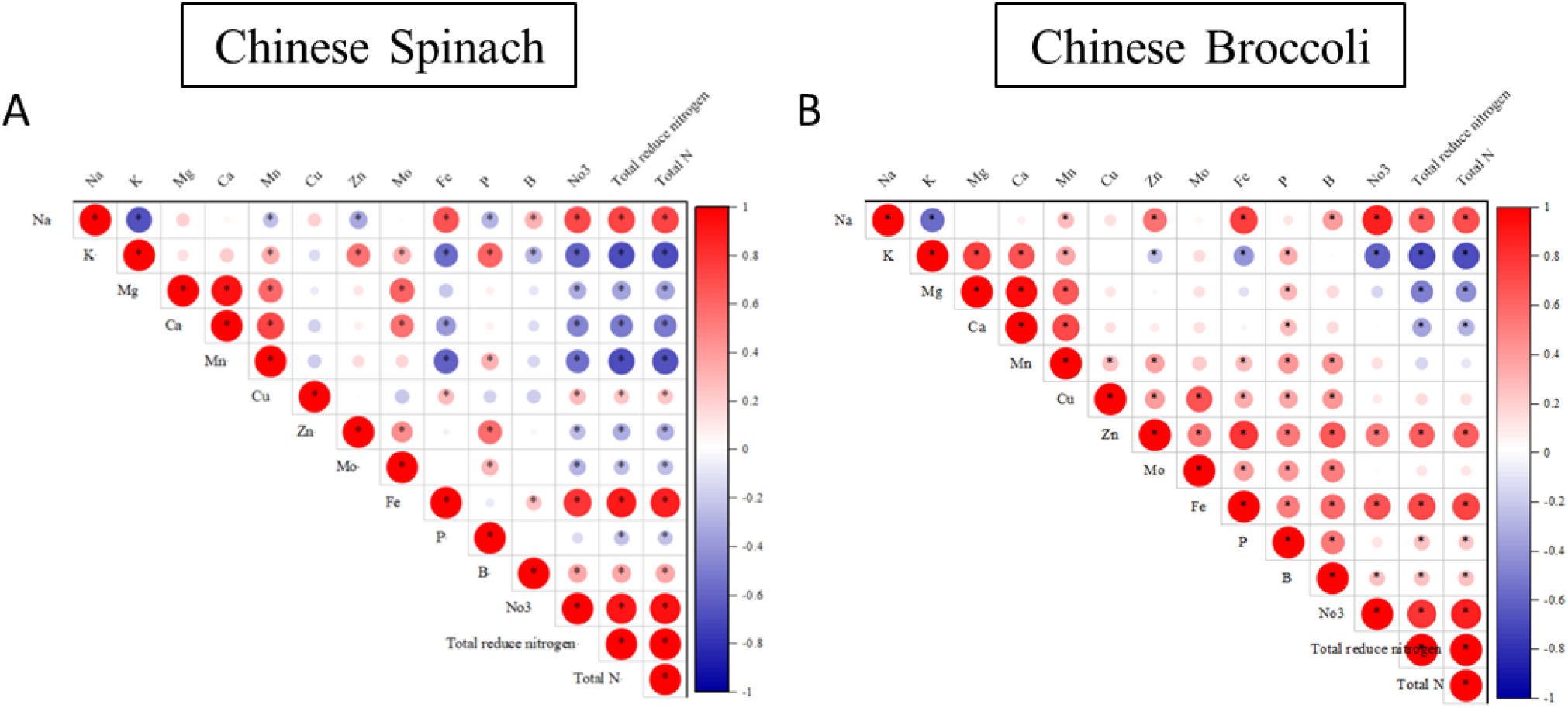
Correlation matrix of leaves mineral concentration. Pearson. linear correlation matrix on harvested leaves mineral concentration on (A) Chinese Spinach and (B) Chinese Broccoli. Colors and circle size represents the Pearson. correlation R2 value according to the color bar. Asterisk represent significance difference using Pearson correlation coefficient (P-value: ≤0.05, 9≤N≤11).

## Discussion

Mineral nutrition balance is essential for growth, development, and disease protection in plants (Hawkesford et al., 2012; Tripathi et al., 2022). As a suggested definition by Arnon, 1950, an essential mineral is not replaceable by other element and has direct or indirect action in plant metabolism. Hence, the ability of the plant to keep mineral homeostasis under changing conditions is crucial to successful crop production. Nitrate and Ammonium uptake is affected by soil PH and soil water content, which results in significant yield reduction (Ruan et al., 2007; Dijkstra and Cheng, 2008). Yet, the effect of lack of or excessive nitrogen on the uptake of other mineral elements is still being determined, especially in terms of practical crop production and quality. In the current study, we clarify the practical impact on physiological and ionome crosstalk dynamics under a wide range of nitrogen supply for leafy greens. Our results reveal the actual optimum response of key physiological parameters as well as on the effect on the PNB and excess nitrogen runoff.

Plant physiological traits has been affected significantly by the nitrogen treatment. Daily transpiration revealed gradual increase in significance between the N treatments throughout the experiments (Fig. 2, A-C and Fig. 3, A-C respectively). This cumulative transpiration response has been reported before, for example, the response of the transpiration of *S. bicolor* (Sorghum) to salinity stress over time (Dewi et al., 2023). Transpiration is known to correlate with crop yield in Chinese Spinach (Liu and Stützel, 2004), and indeed, the agronomic WUE reveals that the 120, 160, and 200 are also more efficient in using the transpired water to increase dry biomass (Fig. 2, D-E and Fig. 3, D-E). Moreover, N supply is related to positive feedback of plant growth, N supply showed to increase chlorophyll a and b (Mu et al., 2016; Mu and Chen, 2021b). Our results support recent studies that suggest that optimal N supply can enhance plant productivity by three leading causes: 1. Regulation of stomatal conductance without impacting the assimilation rate reduces water loss; 2. Increased N investment in the photosynthetic apparatus increases the assimilation rate; 3. There is a moderate increase in the assimilation rate with a slight decrease in stomatal conductance (Plett et al., 2020). Thus, we suggest that under low N supply transpiration is reduced even if water is not limited factor as N limitation prevent the use of photosynthetic produced sugars. Furthermore, plants may invest more N to bioenergetic processes to support a high electron transport rate during the photosynthesis process during N starvation stress (Mu and Chen, 2021b), which might create additional starvation stress on the pre-stress plants. The improvement of NUE_crop_ under low N concentrations (Fig 2, F and Fig. 3, F) suggest the ability of the plant to cope with nitrogen stress by maximizing its physiological traits per N molecules. A similar observation of NUE_crop_ reduction concerning the N supply was found in Rice (*Oryza Sativa*; Nguyen et al., 2014). In opposition to the starvation response, we suggest that the plant minimize the N toxicity by reducing NUE_crop_ to overcome the N abundance toxicity symptoms when transpiration and photosynthesis decreasing, and plant N requirements reduce. Similar NUE response to N delivery had been shown in previous study on *Cannabis Sativa L*. (Saloner and Bernstein, 2020)

Excess N is one of the most negatively impacting elements on soil health in modern agriculture practice (McLauchlan, 2007). Our results indicate that low N concentrations are associated with elevated PNB and reduced excess nitrogen, whereas high nitrogen concentrations correlate with decreased PNB and increased excess nitrogen (Fig. 2, G and Fig. 3, G). Thus using lower nitrogen concentrations in the fertilizers will facilitate a more environmentally sustainable approach, without significantly impacting key physiological traits and yield, and enhancing ionomic crosstalk balance. Furthermore, discovering the different relationships between minerals under stress conditions could reveal a dedicated compensation mechanism within the plant to extend our knowledge of the ionome-physiological processes within the plant. One of the interesting results is the decrease in K absorbance to the plant along the nitrogen treatment (Fig. 4, C-D). We suggest that this N-K trade-off might occur due to the competition between increasing NH4^+^ along with the N concentration treatment increase and K transport to the cells, which might explain our results (White, 1996; Hoopen et al., 2010). P response crosstalk with N was much less substantial than K, where only in the Chinese broccoli some optimum curve reveled around N concentration of 80 ppm (Fig. 4, E-F). Recent studies reported mutual N-P interaction, as N showed to regulate P starvation response suggested by mediating degradation of plasma membrane P transporters (Lin et al., 2013; Medici et al., 2019b), and can suggest N can affect the P concentration by regulating the P transporters. Yet, the exact interaction is still unclear and need further investigation on different crops to understand the relationship. Ca and Mg optimum curve patterns were similar, with moderate positively correlation with N (Fig. 4, I-J, and Fig. 6). Mg has been proven beneficial to growth and yield (Yousaf et al., 2021) and have crucial part in the chlorophyl a molecule (Lincoln Taiz, Eduardo Zeiger, Ian M. Møller, 2015). Fe (Lee et al., 2024) was highly correlated to the N treatment (Fig. 5, A-B; Fig 6; Fig S2). We suggest two mechanisms that might explain that. First, is the fact that Fe is an immobile nutrient in the plant, thus, as fater the plant grow due to ample N conditions the higher the need for new Fe to be absorbed and transformed to the new tissues. Secondly, we suggest that as Fe is a fundamental factor in several Nitrate and ammonium assimilation pathway enzymes (e.g. nitrate reductase, Nitrite reductase, and Glutamate synthase;(Berges and Mulholland, 2008; Heldt and Piechulla, 2011), higher N concentration will result an increase in the nitrogen assimilation pathway, following enzymatic activity increase, and higher demand for Fe within the cell. A cultivar-specific effect was shown on the Zn concentration, whereas the Chinese Broccoli concentration was induced on higher N treatments, while on Chinese Spinach, no significant pattern was revealed. Mn concentration reduces the physiological toxic concentration, which might serve as an indicator for N toxicity in plants.

## Supporting information

Suppelemntary information

## Acknowledgements

I would like to express my deepest appreciation to Tan Li Yi for the crucial help during the research. The study was founded by the National Research Foundation Singapore under its campus for Research Excellence and Technological Enterprise (CREATE) programme.

## Author contribution

I.T played a pivotal role in the planning and formulation of the hypothesis, conducted all the experiments, preformed statistical analysis, managed plant growth and post-harvest analysis and was primary contributed to the writing of this manuscript. A.H was part of the harvest processing. M.R.P.T was part of the harvest processing and consulted on statistical analysis. Z.Z was part of the harvest processing. Q.L was actively involved in the leaf mineral analysis procedure. M.I.S managed fertilizer preparation. D.T supported the plant growth. H.J revised the manuscript. I.H revised the manuscript. K.W revised the manuscript. M.G revised the manuscript. M.M as the principal investigator, managed the experiments, contributed to hypothesis generation, experimental design, data analysis, writing and revising the manuscript.

## Data availability

The data supporting this study’s findings are available from the corresponding authors upon reasonable request. Additionally, some of the data are located in this article’s supplementary material.

## Reference

Berges JA, Mulholland MR (2008) Enzymes and Nitrogen Cycling. Nitrogen in the Marine Environment 1385–1444

Cakmak I, Marschner H (1990) Decrease in nitrate uptake and increase in proton release in zinc deficient cotton, sunflower and buckwheat plants. Plant Soil 129: 261–268

Congreves KA, Otchere O, Ferland D, Farzadfar S, Williams S, Arcand MM (2021) Nitrogen Use Efficiency Definitions of Today and Tomorrow. Front Plant Sci 12: 637108

Dalal A, Shenhar I, Bourstein R, Mayo A, Grunwald Y, Averbuch N, Attia Z, Wallach R, Moshelion M (2020) A Telemetric, Gravimetric Platform for Real-Time Physiological Phenotyping of Plant–Environment Interactions. JoVE (Journal of Visualized Experiments) 2020: e61280

Dewi ES, Abdulai I, Bracho-Mujica G, Appiah M, Rötter RP (2023) Agronomic and Physiological Traits Response of Three Tropical Sorghum (Sorghum bicolor L.) Cultivars to Drought and Salinity. Agronomy 13: 2788

Dijkstra FA, Cheng W (2008) Increased soil moisture content increases plant N uptake and the abundance of 15N in plant biomass. Plant Soil 302: 263–271

Fan X, Zhou X, Chen H, Tang M, Xie X (2021) Cross-Talks Between Macro- and Micronutrient Uptake and Signaling in Plants. Front Plant Sci 12: 663477

Gniazdowska A, Rychter AM (2000) Nitrate uptake by bean (Phaseolus vulgaris L.) roots under phosphate deficiency. Plant Soil 226: 79–85

Hartemink AE, Barrow NJ (2023) Soil pH - nutrient relationships: the diagram. Plant Soil 486: 209–215

Hawkesford M, Horst W, Kichey T, Lambers H, Schjoerring J, Møller IS, White P (2012) Functions of Macronutrients. Marschner’s Mineral Nutrition of Higher Plants: Third Edition 135–189

Heldt H-W, Piechulla B (2011) Nitrate assimilation is essential for the synthesis of organic matter. Plant Biochemistry 273–305

Hoopen F Ten, Cuin TA, Pedas P, Hegelund JN, Shabala S, Schjoerring JK, Jahn TP (2010) Competition between uptake of ammonium and potassium in barley and Arabidopsis roots: molecular mechanisms and physiological consequences. J Exp Bot 61: 2303–2315

Ji C, Li J, Jiang C, Zhang L, Shi L, Xu F, Cai H (2022) Zinc and nitrogen synergistic act on root-to-shoot translocation and preferential distribution in rice. J Adv Res 35: 187–198

Kumar S, Kumar S, Mohapatra T (2021) Interaction Between Macro- and Micro-Nutrients in Plants. Front Plant Sci 12: 665583

Lee HW, Bi X, Henry CJ (2024) Comparative evaluation of minerals content of common green leafy vegetables consumed by the Asian populations in Singapore. Journal of Food Composition and Analysis 125: 105787

Lin WY, Huang TK, Chiou TJ (2013) NITROGEN LIMITATION ADAPTATION, a Target of MicroRNA827, Mediates Degradation of Plasma Membrane–Localized Phosphate Transporters to Maintain Phosphate Homeostasis in Arabidopsis. Plant Cell 25: 4061–4074

Lincoln Taiz, Eduardo Zeiger, Ian M. Møller AM (2015) Plant Physiology (Fourth Edition). Accumulation and Partitioning of Photosynthates—Starch and Sucrose 230–237

Liu F, Stützel H (2004) Biomass partitioning, specific leaf area, and water use efficiency of vegetable amaranth (Amaranthus spp.) in response to drought stress. Sci Hortic 102: 15–27

Maillard A, Etienne P, Diquélou S, Trouverie J, Billard V, Yvin JC, Ourry A (2016) Nutrient deficiencies modify the ionomic composition of plant tissues: a focus on cross-talk between molybdenum and other nutrients in Brassica napus. J Exp Bot 67: 5631–5641

Marschner H (2011) Mineral Nutrition of Higher Plants 3rd Edition eBook ISBN: 9780123849069. eBook ISBN: 9780123849069

McLauchlan K (2007) The Nature and Longevity of Agricultural Impacts on Soil Carbon and Nutrients: A Review. Ecosystems 2007 9:8 9: 1364–1382

Medici A, Marshall-Colon A, Ronzier E, Szponarski W, Wang R, Gojon A, Crawford NM, Ruffel S, Coruzzi GM, Krouk G (2015) AtNIGT1/HRS1 integrates nitrate and phosphate signals at the Arabidopsis root tip. Nature Communications 2015 6:1 6: 1–11

Medici A, Szponarski W, Dangeville P, Safi A, Dissanayake IM, Saenchai C, Emanuel A, Rubio V, Lacombe B, Ruffel S, et al (2019a) Identification of Molecular Integrators Shows that Nitrogen Actively Controls the Phosphate Starvation Response in Plants. Plant Cell 31: 1171– 1184

Medici A, Szponarski W, Dangeville P, Safi A, Dissanayake IM, Saenchai C, Emanuel A, Rubio V, Lacombe B, Ruffel S, et al (2019b) Identification of Molecular Integrators Shows that Nitrogen Actively Controls the Phosphate Starvation Response in Plants. Plant Cell 31: 1171– 1184

Mu X, Chen Q, Chen F, Yuan L, Mi G (2016) Within-leaf nitrogen allocation in adaptation to low nitrogen supply in maize during grain-filling stage. Front Plant Sci 7: 194737

Mu X, Chen Y (2021a) The physiological response of photosynthesis to nitrogen deficiency. Plant Physiology and Biochemistry 158: 76–82

Mu X, Chen Y (2021b) The physiological response of photosynthesis to nitrogen deficiency. Plant Physiology and Biochemistry 158: 76–82

Mu X, Chen Y (2021c) The physiological response of photosynthesis to nitrogen deficiency. Plant Physiology and Biochemistry 158: 76–82

Ngosong C, Bongkisheri V, Tanyi CB, Nanganoa LT, Tening AS (2019) Optimizing Nitrogen Fertilization Regimes for Sustainable Maize (Zea mays L.) Production on the Volcanic Soils of Buea Cameroon. Advances in Agriculture 2019: 4681825

Nguyen HTT, Van Pham C, Bertin P (2014) The effect of nitrogen concentration on nitrogen use efficiency and related parameters in cultivated rices (Oryza sativa L. subsp. indica and japonica and O. glaberrima Steud.) in hydroponics. Euphytica 198: 137–151

Niu J, Chen F, Mi G, Li C, Zhang F (2007) Transpiration, and Nitrogen Uptake and Flow in Two Maize (Zea mays L.) Inbred Lines as Affected by Nitrogen Supply. Ann Bot 99: 153–160

Plett DC, Ranathunge K, Melino VJ, Kuya N, Uga Y, Kronzucker HJ (2020) The intersection of nitrogen nutrition and water use in plants: new paths toward improved crop productivity. J Exp Bot 71: 4452–4468

Ruan J, Jó J, Gerendàs J, Gerendàs G, Härdter R, Härdter H, Sattelmacher B (2007) Effect of Nitrogen Form and Root-zone pH on Growth and Nitrogen Uptake of Tea (Camellia sinensis) Plants. Ann Bot 99: 301–310

Saloner A, Bernstein N (2020) Response of Medical Cannabis (Cannabis sativa L.) to Nitrogen Supply Under Long Photoperiod. Front Plant Sci 11: 572293

Singh H, Northup BK, Rice CW, Prasad PVV (2022) Biochar applications influence soil physical and chemical properties, microbial diversity, and crop productivity: a meta-analysis. Biochar 4: 1–17

Smith FW, Jackson WA (1987) Nitrogen Enhancement of Phosphate Transport in Roots of Zea mays L: II. Kinetic and Inhibitor Studies. Plant Physiol 84: 1319–1324

Tripathi R, Tewari R, Singh KP, Keswani C, Minkina T, Srivastava AK, De Corato U, Sansinenea E (2022) Plant mineral nutrition and disease resistance: A significant linkage for sustainable crop protection. Front Plant Sci 13: 883970

Uhart SA, Andrade FH (1995) Nitrogen Deficiency in Maize: II. Carbon-Nitrogen Interaction Effects on Kernel Number and Grain Yield. Crop Sci 35: 1384–1389

Wang D, Xu Z, Zhao J, Wang Y, Yu Z (2011) Excessive nitrogen application decreases grain yield and increases nitrogen loss in a wheat–soil system. Acta Agric Scand B Soil Plant Sci 61: 681– 692

White PJ (1996) The permeation of ammonium through a voltage-independent K+ channel in the plasma membrane of rye roots. Journal of Membrane Biology 152: 89–99

Williams L, Salt DE (2009) The plant ionome coming into focus. Curr Opin Plant Biol 12: 247

Yu CQ, Huang X, Chen H, Godfray HCJ, Wright JS, Hall JW, Gong P, Ni SQ, Qiao SC, Huang GR, et al (2019) Managing nitrogen to restore water quality in China. Nature 2019 567:7749 567: 516–520

Zhang S, Zhao Y, Shi R, Waterhouse GIN, Zhang T (2019) Photocatalytic ammonia synthesis: Recent progress and future. EnergyChem 1: 100013

Zhao X, Zhou Y, Min J, Wang S, Shi W, Xing G (2012) Nitrogen runoff dominates water nitrogen pollution from rice-wheat rotation in the Taihu Lake region of China. Agric Ecosyst Environ 156: 1–11

