## Supplementary material for "Nitrogen ionome dynamics on leafy vegetables in tropical climate": Suppelemntary information

### Slide 1
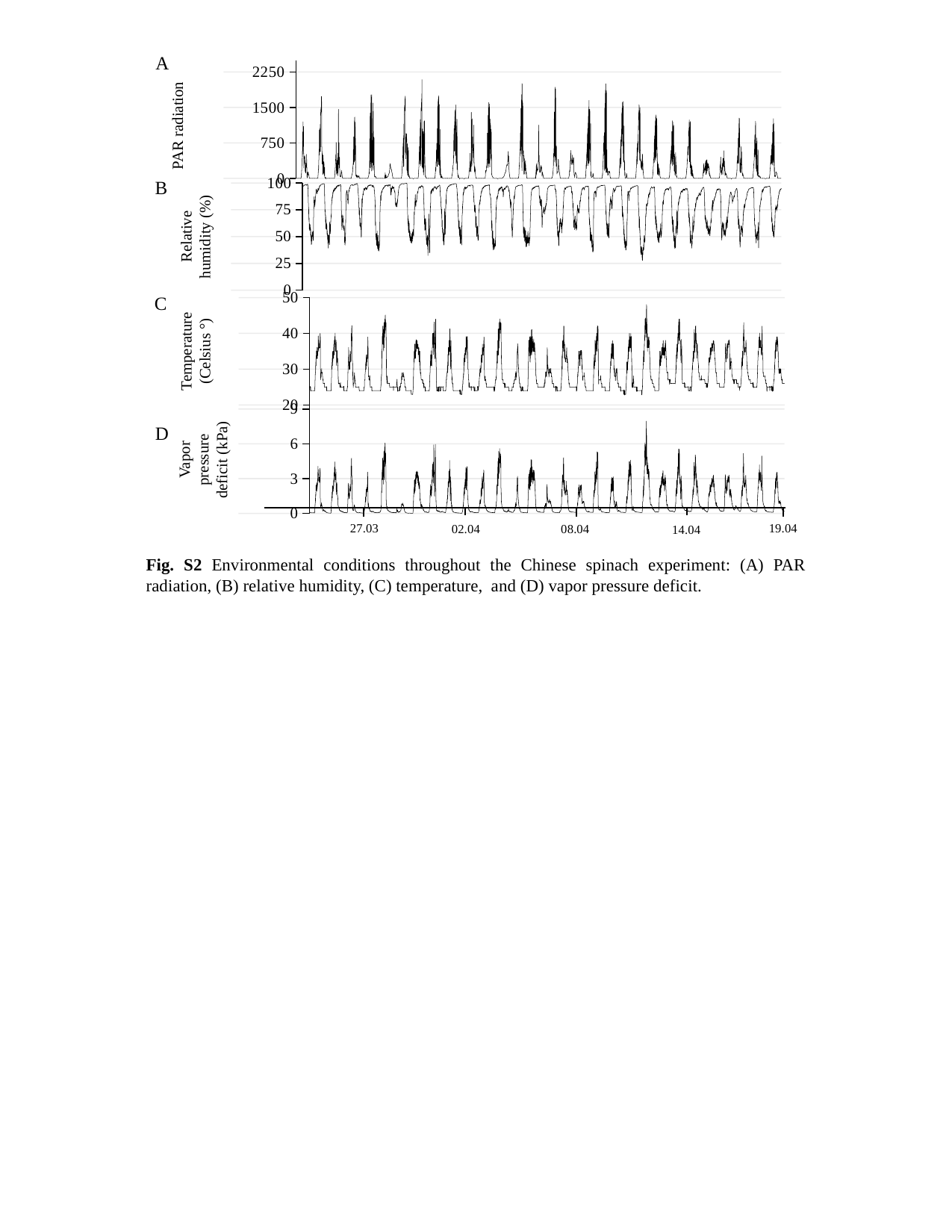

A
#### Chart
| Category | |
|---|---|B
#### Chart
| Category | |
|---|---|Relative humidity (%)
C
#### Chart
| Category | |
|---|---|Temperature (Celsius °)
#### Chart
| Category | |
|---|---|D
Vapor pressure deficit (kPa)
27.03
19.04
08.04
02.04
14.04
Fig. S2 Environmental conditions throughout the Chinese spinach experiment: (A) PAR radiation, (B) relative humidity, (C) temperature, and (D) vapor pressure deficit.

### Slide 2
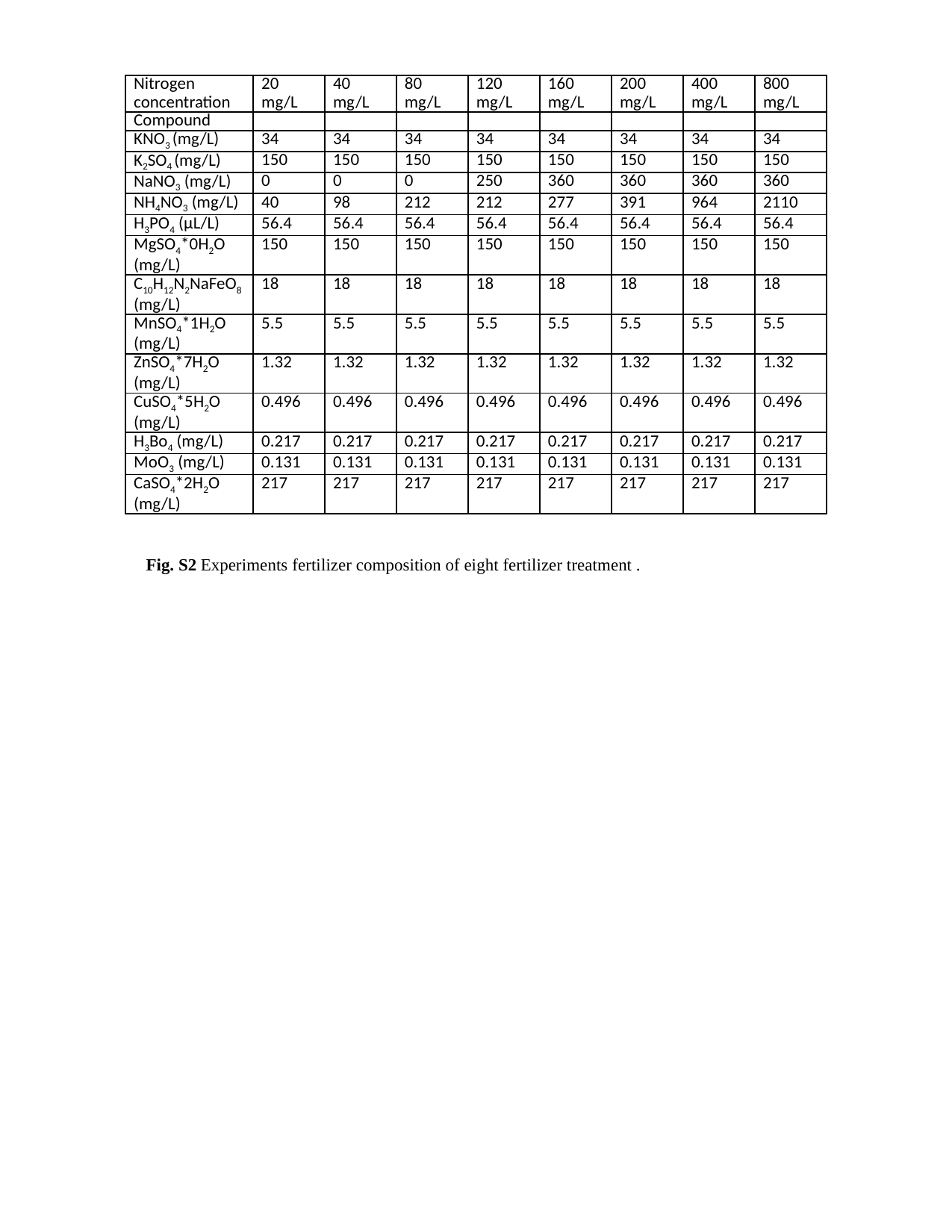

| Nitrogen concentration | 20 mg/L | 40 mg/L | 80 mg/L | 120 mg/L | 160 mg/L | 200 mg/L | 400 mg/L | 800 mg/L |
| --- | --- | --- | --- | --- | --- | --- | --- | --- |
| Compound | | | | | | | | |
| KNO_3_ (mg/L) | 34 | 34 | 34 | 34 | 34 | 34 | 34 | 34 |
| K_2_SO_4_ (mg/L) | 150 | 150 | 150 | 150 | 150 | 150 | 150 | 150 |
| NaNO_3_ (mg/L) | 0 | 0 | 0 | 250 | 360 | 360 | 360 | 360 |
| NH_4_NO_3_ (mg/L) | 40 | 98 | 212 | 212 | 277 | 391 | 964 | 2110 |
| H_3_PO_4_ (µL/L) | 56.4 | 56.4 | 56.4 | 56.4 | 56.4 | 56.4 | 56.4 | 56.4 |
| MgSO_4_\*0H_2_O (mg/L) | 150 | 150 | 150 | 150 | 150 | 150 | 150 | 150 |
| C_10_H_12_N_2_NaFeO_8_ (mg/L) | 18 | 18 | 18 | 18 | 18 | 18 | 18 | 18 |
| MnSO_4_\*1H_2_O (mg/L) | 5.5 | 5.5 | 5.5 | 5.5 | 5.5 | 5.5 | 5.5 | 5.5 |
| ZnSO_4_\*7H_2_O (mg/L) | 1.32 | 1.32 | 1.32 | 1.32 | 1.32 | 1.32 | 1.32 | 1.32 |
| CuSO_4_\*5H_2_O (mg/L) | 0.496 | 0.496 | 0.496 | 0.496 | 0.496 | 0.496 | 0.496 | 0.496 |
| H_3_Bo_4_ (mg/L) | 0.217 | 0.217 | 0.217 | 0.217 | 0.217 | 0.217 | 0.217 | 0.217 |
| MoO_3_ (mg/L) | 0.131 | 0.131 | 0.131 | 0.131 | 0.131 | 0.131 | 0.131 | 0.131 |
| CaSO_4_\*2H_2_O (mg/L) | 217 | 217 | 217 | 217 | 217 | 217 | 217 | 217 |
Fig. S2 Experiments fertilizer composition of eight fertilizer treatment .
